# Defining the minimal enzymatic requirements for fatty acid scavenging from lysophosphatidylcholine by erythrocytic *Plasmodium falciparum*

**DOI:** 10.1101/2025.08.07.669178

**Authors:** Jiapeng Liu, Katherine R. Fike, Seema Dalal, Christie Dapper, Michael Klemba

**Affiliations:** Department of Biochemistry, Virginia Tech, Blacksburg, VA 24061, USA

**Keywords:** malaria, lysophosphatidylcholine, lysophospholipase, fatty acid, choline, metabolism

## Abstract

Host-derived lysophosphatidylcholine (LPC) is a significant source of choline and fatty acids for the intraerythrocytic malaria parasite *Plasmodium falciparum*. Two lysophospholipases play a dominant role in LPC catabolism: exported lipase 2 (XL2) and exported lipase homolog 4 (XLH4). Loss of these two enzymes greatly reduces, but does not abrogate, the parasite’s ability to utilize LPC as a source of fatty acids. In this study, we identify a third enzyme, termed “prodrug activation and resistance esterase” (PARE), that mediates low levels of LPC hydrolysis. Loss of PARE alone had no effect on the parasite’s ability to scavenge fatty acids from LPC. However, when combined with the loss of XL2 and XLH4, knockdown of PARE impacted the parasite’s ability to scavenge both choline and fatty acids from LPC. Furthermore, PARE/XL2/XLH4-deficient parasites were unable to complete a replication cycle when cultured in defined media with LPC as the sole source of exogenous fatty acids. We show that PARE is a membrane-associated enzyme with a substantial presence at the parasite periphery and propose a model whereby PARE catalyzes the hydrolysis of inwardly-diffusing LPC. Our findings reveal that asexual *P. falciparum* is dependent on parasite-encoded enzymes for LPC catabolism and rule out host erythrocyte enzymes as a physiologically-relevant source of lysophospholipase activity.

## 1. INTRODUCTION

The human malaria parasite *Plasmodium falciparum* relies on exogenous sources of fatty acids as it undergoes asexual replication within the host erythrocyte. A copious supply of fatty acids is required to support vigorous phospholipid and neutral lipid synthesis during the 48-hour replication cycle, which produces up to 32 daughter parasites (Vial and Ancelin, 1998). While the parasite genome encodes a fatty acid synthase, it can be knocked out in blood-stage parasites and is not considered to be a significant source of fatty acids during asexual replication (Vaughan et al., 2009; Yu et al., 2008). There are two major exogenous sources of fatty acids in the host circulation: unesterified or free fatty acids, which are present in serum at concentrations of ∼200 µM, and esterified fatty acids in the form of lysophosphatidylcholine (LPC), which are present at similar concentrations (Psychogios et al., 2011). Metabolic labeling experiments have revealed that both free fatty acids and those derived from LPC can be incorporated into *P. falciparum* lipids (Brancucci et al., 2017; Vial et al., 1989; Vial et al., 1982), demonstrating a flexibility with respect to fatty acid acquisition. Parasites can complete a replication cycle in media containing as few as two free fatty acids or LPC species, provided that one each of saturated and unsaturated fatty acid/acyl groups is present (Liu et al., 2024; Mi-Ichi et al., 2007). In addition to supplying fatty acids, LPC provides a critical source of choline for parasite phosphatidylcholine synthesis *via* the Kennedy pathway (Brancucci et al., 2017; Ramaprasad et al., 2022; Wein et al., 2018). Interestingly, depletion of exogenous LPC serves as an environmental cue for gametocytogenesis, a phenomenon that appears to be related to its provision of choline (Brancucci et al., 2017).

The enzymology of LPC catabolism in *P. falciparum*-infected erythrocytes has recently come into focus. We have identified two enzymes, termed exported lipase 2 (XL2) and exported lipase homolog 4 (XL4) that are responsible for over 90% of *P. falciparum*-encoded lysophospholipase activity (Liu et al., 2024). XL2 is exported to the host erythrocyte, whereas XLH4 is retained within the parasite. Both are α/μ hydrolase-class serine hydrolases that exhibit A-type lysophospholipase activity, yielding a fatty acid and glycerophosphocholine (GPC) as the LPC hydrolysis products. Disruption of both XL2 and XLH4 reduced the parasite’s ability to scavenge fatty acids from LPC and sensitized parasites to LPC toxicity (Liu et al., 2024).

Hydrolysis of GPC to choline and glycerol-3-phosphate is catalyzed by a parasite-encoded glycerophosphodiester phosphodiesterase (GDPD), which is located in the parasitophorous vacuole (PV) and cytosol and is therefore poised to act on GPC generated by both XL2 and XLH4 (Denloye et al., 2012; Ramaprasad et al., 2022). Conditional knockout of GDPD revealed that the enzyme is solely responsible for the release of choline from LPC-derived GPC and that this an essential function under standard culture conditions (Ramaprasad et al., 2022). Thus, the parasite’s ability to efficiently metabolize LPC may be driven by its requirement for choline.

While XL2 and XLH4 constitute the dominant lysophospholipase activities in parasitized erythrocytes, disruption of both reduced, but did not abolish, the ability of parasites to replicate in a defined culture medium consisting of 16:0 and 18:1 LPC as the sole sources of exogenous fatty acids (Liu et al., 2024). One possible explanation for this finding is the presence of additional parasite-encoded lysophospholipase activities that, while not sufficient to fully compensate for the loss of XL2 and XLH4, are able to provide enough fatty acids to support a lower replication efficiency. Several other parasite serine hydrolases, termed LPL1, LPL3 and LPL20, have been shown to catalyze LPC hydrolysis *in vitro* (Asad et al., 2021; Sheokand et al., 2021; Sheokand et al., 2023) and are possible candidates; however, their contributions to LPC hydrolysis *in vivo* have not been defined. Alternately, LPC hydrolysis could be catalyzed by a host-derived lysophospholipase activity that has been detected in human erythrocytes (Podolski et al., 1983; Selle et al., 1993; Tamura et al., 1985).

In this study, we sought to identify the source of the residual lysophospholipase activity in erythrocytes infected with XL2/XLH4-deficient parasites and thereby to more completely define the cohort of enzymes responsible for scavenging fatty acids from LPC. We considered the parasite-encoded, α/μ hydrolase-family enzyme “prodrug activation and resistance esterase” (PARE) to be a leading candidate, as prior activity-based profiling studies have revealed that PARE is an abundant serine hydrolase that is inhibited by the pan-lysophospholipase inhibitor AKU-010 (Liu et al., 2024). PARE was first characterized as a catalyst of pepstatin ester hydrolysis and its inactivation resulted in parasite resistance toward this compound (Istvan et al., 2017). A similar resistance-conferring activity has since been demonstrated for two unrelated ester-containing compounds (Butler et al., 2020; Sindhe et al., 2020), which suggests that PARE exhibits a promiscuous esterase activity. It is potently inhibited by monoacylglycerol lipase inhibitors (Elahi et al., 2019) and the lipid-like natural product Salinipostin A (Yoo et al., 2020), which suggests that it may have a lipid substrate, although its physiological substrate and role in the asexual parasite are currently unknown.

To test whether PARE contributes to LPC catabolism in XL/XLH-deficient parasites, we modified the PARE allele in a previously-described quadruple knockout line (QKO: ΔXL1/XL2/XLH3/XLH4; (Liu et al., 2024)) in order to enable a conditional knockdown approach. The effect of PARE knockdown on *in situ* LPC metabolism in *P. falciparum*-infected erythrocytes and on parasite growth on LPC as a sole source of exogenous fatty acids was assessed. In parallel, the effect of a functional PARE knockout on a wild-type background was examined to determine the effect of a PARE deletion in the presence of XL/XLH enzymes. The subcellular distribution of PARE was determined to provide a spatial context for its contribution to LPC hydrolysis.

## 2. MATERIALS AND METHODS

### 2.1 Reagents

BODIPY™ 500/510 C_1_, C_12_ (4,4-difluoro-5-methyl-4-bora-3a,4a-diaza-*s*-indacene-3-dodecanoic acid), BODIPY-TR-ceramide (BTC) and fatty acid-free BSA were from ThermoFisher. LPC 18:1 and 16:0, (*Z*)-octadec-9-en-17-ynoic acid (oleic acid alkyne), and palmitic and oleic acids were obtained from Avanti Polar Lipids. 3-azido-7-hydroxycoumarin was acquired from Abcam. IDFP, JW642, anhydrotetracycline hydrochloride and blasticidin S hydrochloride were purchased from Cayman Chemical.

### 2.2 Parasite culture

*P. falciparum* clone 3D7 and the knockout lines 3D7-R4 (carrying a nonsense mutation at codon 139 of the PARE coding sequence (Istvan et al., 2017) and referred to here as "ΔPARE") and ΔXL1/2/XLH3/4 (referred to here as “quadruple knockout” or QKO; (Liu et al., 2024)) were cultured in human O^+^ erythrocytes (Grifols Bio Supplies, Inc) at 2% hematocrit in RPMI 1640 medium supplemented with 0.37 mM hypoxanthine, 11 mM glucose, 27 mM sodium bicarbonate, 10 μg/mL gentamicin and 5 g/L Albumax I (Gibco), unless otherwise stated. Cultures were incubated at 37 °C in a 5% CO_2_ with ambient O_2_ and were synchronized by treatment with 5% (w/v) sorbitol.

### 2.3 Generation of transgenic parasite lines

To generate the PARE conditional knockdown line: installation of a 3x hemagglutinin (HA) tag and a TetR aptamer array at the 3’ end of the PARE coding sequence (PF3D7_0709700) in the QKO parasite line was achieved by CRISPR/Cas9 editing (**Fig. 1A**) (Rajaram et al., 2020). A Cas9 sgRNA sequence homologous to bases 1085-1104 at the 3’ end of the PARE coding sequence was inserted into the BtgZ1 site of pUF-Cas9-pre-sgRNA (Garten et al., 2018) (oligonucleotide sequences referenced in this section are provided in **Supplementary Table 1**). For the homology repair plasmid, two homology arm sequences corresponding to ∼350 nucleotides of the PARE coding and 3’ UTR sequences were PCR amplified using primers 1387/1388 and 1389/1390, respectively. Three shield mutations were introduced to prevent Cas9 cleavage of the repair sequence. The two PCR products were merged by overlapping PCR and the product was inserted into AscI*-* and AatII*-* digested pKD^PfAUBL^ (Rajaram et al., 2020), generating pKD-PARE. 75 µg each of pUF-Cas9-pre-sgRNA plasmid and pKD-PARE (linearized with EcoRV) were co-transfected into ring-stage QKO parasites, which were cultured with 0.5 μM anhydrotetracycline (aTc) to maintain PARE expression. After 48 hours, parasites were cultured in medium containing 1.5 μM DSM-1 and 2.5 μg/ml blasticidin S for 7 days.

**Figure 1:**
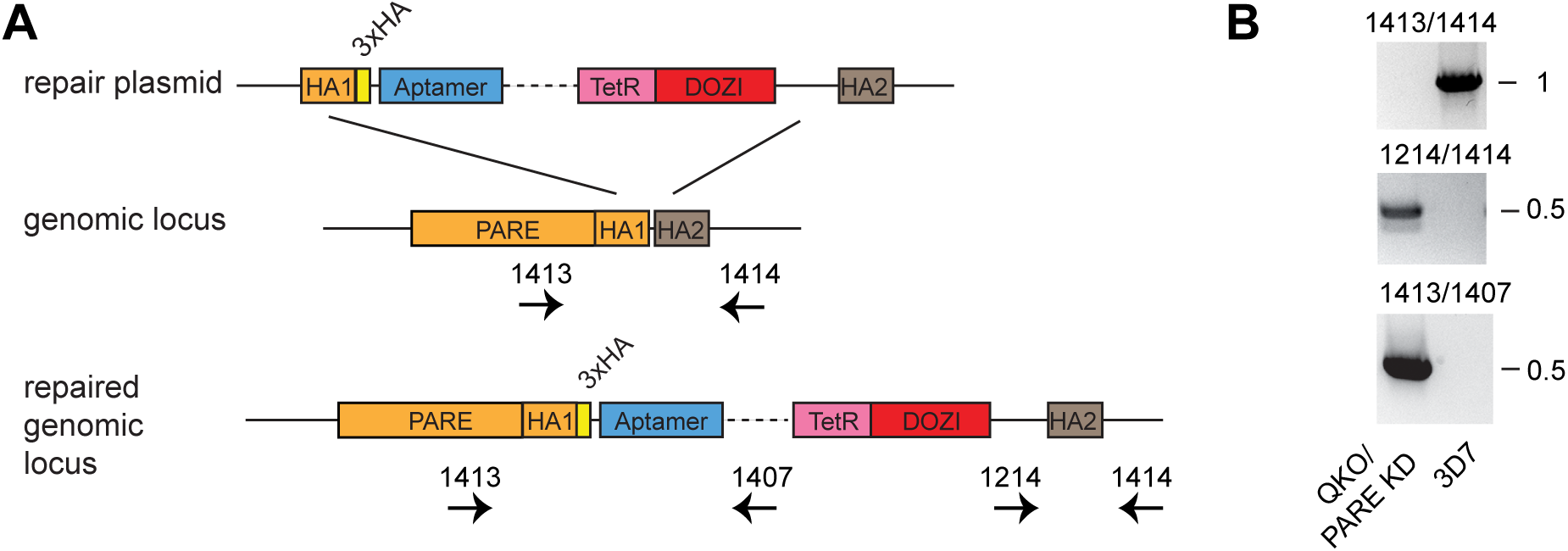
Modification of the PARE allele to introduce TetR-DOZI-mediated conditional knockdown and a C-terminal 3xHA tag. (**A**) Schematic diagram of CRISPR editing. Protein coding sequence is indicated with orange and 3’ UTR sequence with brown. HA, homology arm; 3xHA, triple hemagglutinin tag. Positions of oligos used for diagnostic PCR are indicated. Diagram not drawn to scale. (**B**) PCR genotyping of QKO/PARE KD and wild-type 3D7 genomic DNA confirms the expected modification. Sizes of markers are indicated at right in kilobases.

Parasites emerging from selection were cloned by limiting dilution; clone 8 was used in these studies. Editing at the PARE locus was confirmed by PCR (**Fig. 1B**). The QKO/PARE knockdown (KD) line was routinely cultured in the presence of 0.5 µM aTc. Generation of a parasite line expressing a fusion of the YFP allele Citrine to the C-terminus of endogenous PARE has been described previously (Elahi et al., 2019).

### 2.4 Characterization of PARE expression in knockout and knockdown lines

Pellets of saponin-isolated parasites were resuspended in cold PBS containing 5 μM pepstatin A and 10 µM E-64 to a density of 5 x 10^8^ parasites/mL. The parasite suspension was sonicated three times for eight seconds using a microtip at 70% maximum power and was clarified by centrifugation at 12,000 x *g* at 4 °C. Supernatants were aliquoted, snap frozen in liquid nitrogen and stored at – 80 °C. TAMRA-FP labeling reactions were carried out with 1 µM probe as previously described (Liu et al., 2024). For separation of soluble and membrane fractions, parasite lysate was treated with 1 µM TAMRA-FP and then subjected to ultracentrifugation at 100K x *g* for 30 minutes. The soluble fraction was removed, while the membrane fraction was rinsed gently and then dissolved in 1x denaturing SDS sample buffer.

Proteins were resolved on a 10% polyacrylamide gel and were imaged on a Typhoon RGB flatbed scanner using the 532 nm laser and a 570/20 bandpass filter.

For immunoblot analysis of PARE-3xHA, lysates of saponin-isolated cultures of QKO/PARE KD grown without aTc for a minimum of 10 days, or with aTc, were labeled with the fluorescent reagent QuickStain (Cytiva). The reaction was quenched by adding SDS sample buffer containing 1.25 mM lysine. Proteins were resolved on a 12% SDS-polyacrylamide gel and QuickStain in-gel fluorescence was imaged on a Typhoon RGB flatbed scanner. Proteins were then transferred to a nitrocellulose membrane. The blot was developed with a rabbit polyclonal anti-HA antibody (Invitrogen 71-5500) at 0.1 µg/mL and an HRP-conjugated secondary antibody at 1 ng/mL.

### 2.5 Growth assays

To assess the effect of choline on growth rate, synchronized ring-stage parasites (0-16 h post-invasion) were cultured in complete RPMI (which contains 21 µM choline), or in complete RPMI supplemented with an additional 2 mM choline chloride. Parasites were seeded at 0.7% parasitemia and 2% hematocrit. Every two days over an eight-day period, parasitemia was counted from Giemsa-stained smears (minimum of 1000 cells) and parasites were sub-cultured to 0.7%. “Cumulative parasitemia” was calculated as the product of fold-change in parasitemia and was natural-log transformed. Data were fit by linear regression using Graphpad Prism 10.

### 2.6 Replication in media with defined fatty acid sources

Single-cycle replication of parasites in media with defined exogenous fatty acid sources (LPC 16:0 and 18:1 at various concentrations, or the corresponding two free fatty acids at 30 µM each) were conducted in incomplete RPMI supplemented with 3 mg/mL fatty-acid free BSA as previously described (Liu et al., 2024). Briefly, synchronized ring-stage parasites were transferred to the respective media and cultured at 37 °C in a 5% CO_2_ incubator for 60 hours.

New ring-stage parasites were counted from Giemsa-stained smears (minimum of 1000 cells).

To follow development at various points during the replication cycle, parasites were tightly synchronized by inhibiting egress with the cGMP-dependent protein kinase inhibitor ML10 (Ressurreicao et al., 2020) to accumulate mature schizonts, followed by ML10 washout and, after sufficient reinvasion, sorbitol treatment, yielding a ∼4 hour invasion window. Cultures were sampled at 53 h post-invasion for preparation of Giemsa-stained smears.

### 2.7 Assays for inhibition of in situ LPC hydrolysis

LPC competition assays employing oleic acid alkyne or a fluorescent fatty acid analog were conducted as previously described (Dapper et al., 2022; Liu et al., 2024) with the modification that BODIPY™ 500/510 C_1_, C_12_ was substituted for BODIPY™ 500/510 C_4_, C_9_, the latter being no longer commercially available. Briefly, parasites were labeled with 1 µM BODIPY-TR-ceramide in complete RPMI for 1 hour at 37 °C in a 5% CO_2_ incubator. Parasites were then washed with fatty acid-free RPMI medium for 3 times and incubated with pre-warmed incomplete RPMI containing 3 mg/mL fatty acid-free BSA, 30 µM probe (oleic acid alkyne or BODIPY™ 500/510 C_1_, C_12_) and 30 µM LPC 18:1 (or 50% ethanol for no-LPC controls) for 40 minutes with gentle mixing on an orbital rotator at 37 °C in a CO_2_ incubator. Reactions were harvested and lipids were extracted and analyzed by thin-layer chromatography as previously described (Dapper et al., 2022; Liu et al., 2024).

*2.8 Live-cell fluorescence imaging of PARE-YFP*

Live infected erythrocytes were imaged on an epifluorescence AxioImager M1 microscope equipped with a 100x/NA1.4 objective and an Axiocam MRm camera. Nucleic acid was stained with 0.13 µM Hoechst 33342. Saponin-isolated parasites were obtained by resuspending a parasite culture in complete RPMI supplemented with 40 µg/mL saponin for 10 minutes at room temperature. Parasites were obtained by centrifugation at 1,940 x g, were taken up in RPMI, and were imaged immediately. Image contrast was adjusted using Adobe Photoshop 2025.

## 3. RESULTS

### 3.1 Characterization of PARE-depleted lines

To determine whether PARE is responsible for the residual LPC hydrolysis observed in QKO (ΔXL1/XL2/XLH3/XLH4) parasites, we generated a PARE conditional knockdown line on the QKO background using the tetracycline repressor (TetR)-"development of zygote inhibited" (DOZI) system (Ganesan et al., 2016; Rajaram et al., 2020). An array of TetR-binding aptamers was installed at the 3’ UTR of the PARE gene and a triple-hemagglutinin (3xHA) tag was fused to the PARE C-terminus (**Fig. 1**). This line is referred to hereafter as "QKO/PARE KD". When added to the culture medium, the TetR ligand anhydrotetracycline (aTc) prevents TetR from binding the aptamer array and is therefore permissive for PARE translation. In the absence of aTc, the TetR-DOZI fusion binds to the aptamer array and sequesters the PARE mRNA in a translationally-inactive complex, resulting in knockdown at the protein level.

To assess the efficiency of knockdown, the QKO/PARE KD line was cultured for over 3 generations in the presence and absence of aTc. PARE levels in lysates were determined using the serine hydrolase-directed, activity-based probe TAMRA-fluorophosphonate (TAMRA-FP (Patricelli et al., 2001); **Fig. 2A**), which we have used previously to characterize the activity of this enzyme (Elahi et al., 2019). Reactions were conducted without and with AKU-010, a pan-lysophospholipase inhibitor (Liu et al., 2024). PARE knockdown appeared to be highly efficient; however, quantitation was confounded by the presence of unrelated, labeled polypeptides in close proximity to PARE. As an alternative, PARE-3xHA levels were determined by immunoblotting, which confirmed highly efficient knockdown (**Fig. 2B**). Comparing relative band intensities, the level of PARE in the absence of aTc is estimated to be <1% of that in its presence. To assess the effects of the loss of PARE on a wild-type background, we used a previously-reported parasite line (3D7-R4; termed ΔPARE here) carrying a nonsense mutation in the PARE coding sequence (Istvan et al., 2017). The absence of active PARE from this line was confirmed by TAMRA-FP labeling of ΔPARE lysate (**Fig. 2C**).

**Figure 2:**
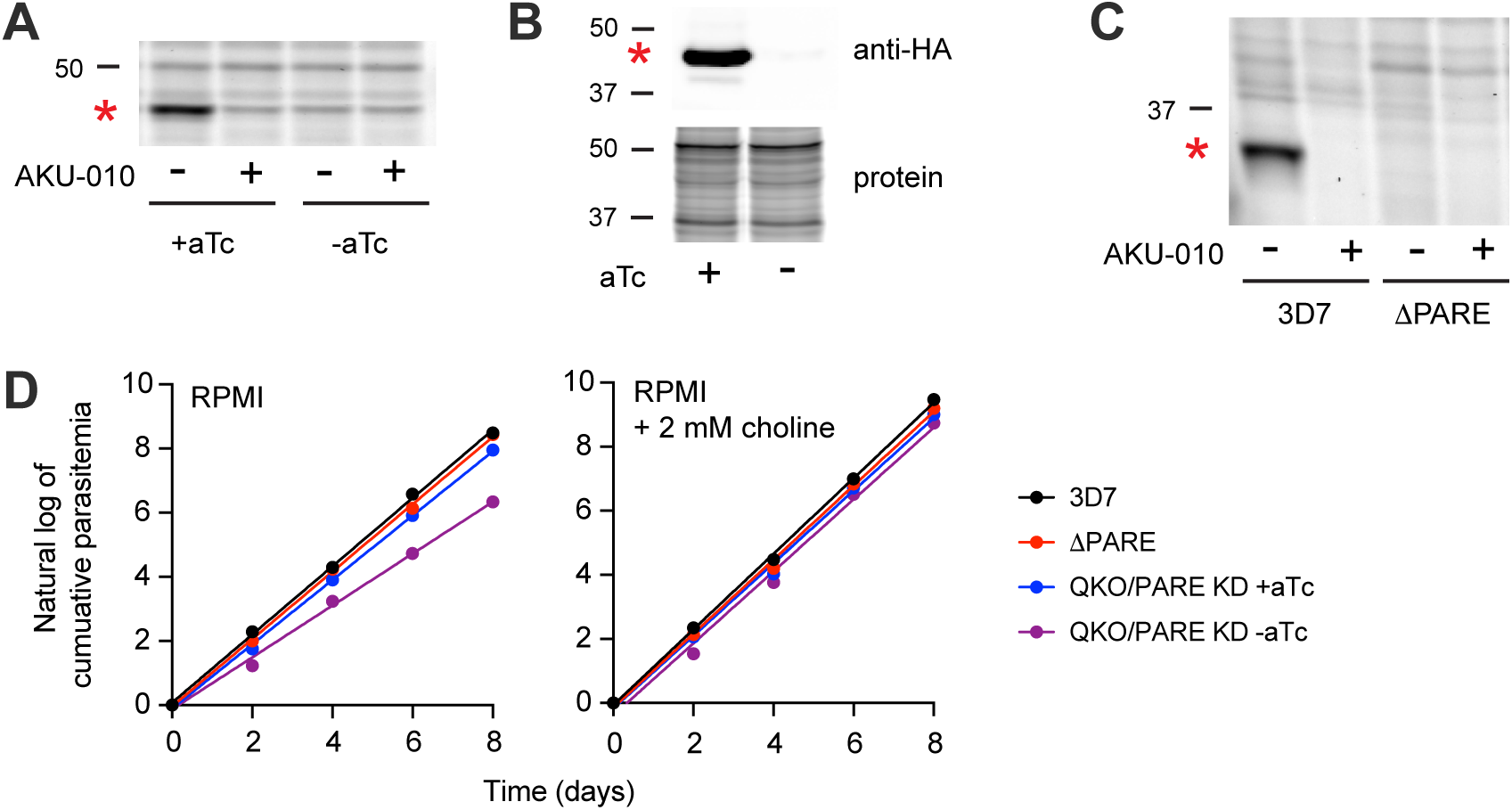
Characterization of PARE-deficient parasite lines. (**A**) TAMRA-FP labeling of PARE in lysates of QKO/PARE KD parasites cultured with and without aTc. The inhibitor AKU-010 (10 µM) was used to determine background labeling. (**B**) Upper panel: Immunolabeling of PARE-3xHA in lysates of QKO/PARE KD parasites cultured with and without aTc. Lower panel: QuickStain protein labeling. (**C**) TAMRA-FP labeling of PARE in lysates of 3D7 and ΔPARE parasites. The inhibitor AKU-010 (10 µM) was used to determine background labeling. In A-C, the position of PARE is indicated with a red asterisk. Sizes of markers are indicated in kDa. Note that the 3xHA tag shifts the apparent molecular mass of tagged PARE (A, B) above that observed for the untagged protein (C). (**D**) Growth of 3D7 and PARE-deficient parasite lines in standard RPMI medium (left panel) or in RPMI supplemented with 2 mM choline (right panel). Results are representative of two independent experiments.

Quantitation of parasite replication rates revealed that the QKO/PARE KD line grew more slowly in the absence of aTc than in its presence (**Fig. 2D**). We speculated that this was due to choline deficiency in the QKO/PARE KD parasites, as LPC has been established as a critical source of exogenous choline (Brancucci et al., 2017; Ramaprasad et al., 2022; Wein et al., 2018) and because free fatty acids are available to the parasite from the serum replacement Albumax (Garcia-Gonzalo and Izpisua Belmonte, 2008). To determine whether choline deficiency was responsible for the observed growth defect, we examined replication rates in RPMI supplemented with an additional 2 mM choline (a concentration that is sufficient to complement a GDPD knockout line (Ramaprasad et al., 2022)). Supplementation restored the growth rate of QKO/PARE KD line (-aTc) to that in the presence of aTc, both of which were essentially identical to those of wild-type and ΔPARE parasites (**Fig. 2D**). Thus, PARE appears to contribute to choline acquisition in the absence of functional XL/XLH enzymes, implying an ability to catalyze LPC hydrolysis.

### 3.2 Loss of PARE reduces in situ LPC hydrolysis in XL/XLH-deficient parasites

To directly evaluate the contribution of PARE to LPC hydrolysis in intact, parasitized erythrocytes (referred to as “*in situ*” hydrolysis), we used fatty acid probe-based LPC competition assays that were previously developed to quantify the contributions of the XL/XLH enzymes (Dapper et al., 2022; Liu et al., 2024). In one version of this assay, unlabeled fatty acids that are liberated from LPC compete with the fatty acid probe oleate alkyne (OA) for incorporation into parasite phospholipids and neutral lipids, leading to reduced levels of alkyne-labeled parasite lipids. We employed this assay to assess the consequences of the loss of PARE activity alone. ΔPARE and wild-type parasites exhibited identical levels of OA incorporation into parasite phospholipids and neutral lipids (**Fig. 3A**), which indicates that the loss of PARE does not substantially reduce the magnitude of *in situ* LPC hydrolysis in the presence of XL2 and XLH4.

**Figure 3:**
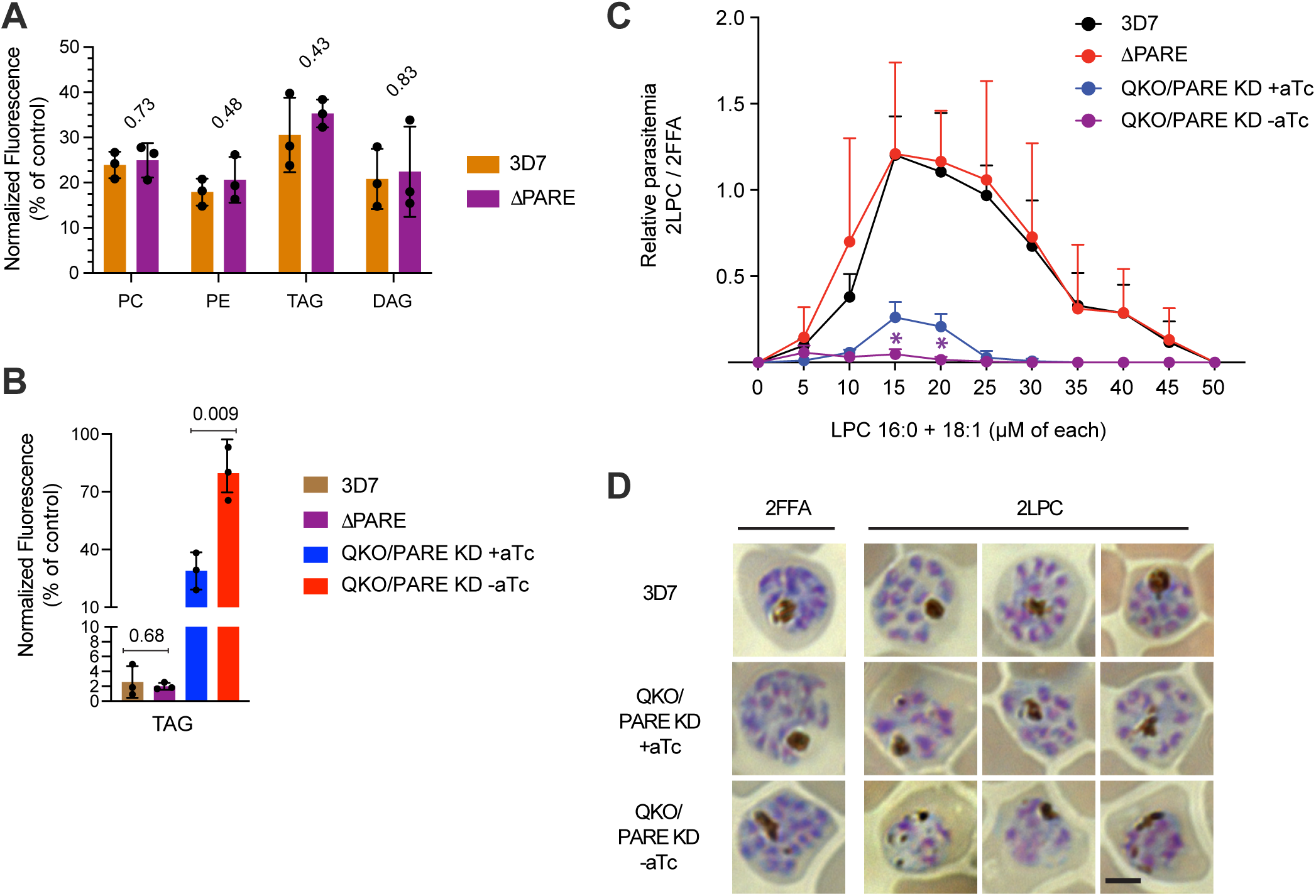
Effect of PARE depletion on fatty acid scavenging from LPC. (**A**) LPC competition assay with 3D7 and ΔPARE parasites using an oleate alkyne probe. (**B**) LPC competition assay with 3D7, ΔPARE, and QKO/PARE KD with and without aTc using the fluorescent fatty acid probe BODIPY 500/510 C_1_, C_12_. In A and B, means and standard deviations are from three independent experiments. The indicated *p*-values were determined using an unpaired, two-tailed Welch’s *t*-test. (**C**) Replication of parasites in media with 16:0 and 18:1 LPC as the sole source of exogenous fatty acids. Ring-stage parasites were cultured in media containing the indicated concentrations of LPC. After 60 h, reinvaded (ring) parasites were counted and the values normalized to those observed in a medium containing two free fatty acids. Means and standard deviations are shown from three independent experiments. Asterisks indicate a *p*-value of <0.05 for an unpaired, two-tailed Welch’s *t*-test for differences between QKO/PARE KD +aTc and - aTc. (**D**) Images of Giemsa-stained parasites that were cultured for 53 h in 2LPC or 2FFA media. Scale bar, 3 µm.

To investigate the effects of PARE depletion on a QKO background, we required a more sensitive assay than that afforded by the OA uptake assay, as the loss of XL2 and XLH4 essentially abrogated the ability of LPC-derived fatty acids to compete with OA incorporation (Liu et al., 2024). We have found that fluorescent fatty acid analogs such as 4,4-difluoro-5-methyl-4-bora-3a,4a-diaza-*s*-indacene-3-dodecanoic acid (tradename BODIPY 500/510 C_1_, C_12_) provide a substantially more sensitive assay, a phenomenon that presumably originates in its reduced ability to compete with natural fatty acids for the active sites of acyl-CoA synthetases and lipid biosynthetic enzymes (Dapper et al., 2022). Even in this sensitive assay, the loss of PARE alone had no effect on the ability of LPC to compete with probe labeling of the neutral lipid triacylglycerol (ΔPARE, **Fig. 3B**). In contrast, the QKO/PARE KD line cultured with aTc exhibited a dramatic increase in label incorporation, as expected for the sharp reduction in exogenous LPC hydrolysis in the absence of XL2 and XLH4. Knockdown of PARE on the QKO background (-aTc) resulted in an additional two-fold increase in fractional probe incorporation, indicating a further reduction in the level of LPC hydrolysis. These experiments reveal that PARE contributes to the *in situ* liberation of free fatty acids from LPC; however, this activity is only observable in an XL/XLH-deficient background.

### 3.3 Loss of PARE abrogates QKO parasite growth when LPC is the sole source of exogenous fatty acids

We next evaluated the consequences of the loss of PARE on wild-type and QKO backgrounds for the ability to use exogenous LPC as the sole source of fatty acids. Ring-stage parasites were cultured in defined media containing either two LPC species (16:0 and 18:1, at varying concentrations; 2LPC media), or the corresponding free fatty acids (at the optimum concentration of 30 µM each; 2FFA medium) to control for the effects of limiting the parasite to two fatty acids. After completion of one replication cycle, re-invaded parasitemias in 2LPC media were counted and normalized to those in 2FFA. We have previously employed this assay to characterize XL/XLH-deficient parasites and found that they have a diminished capacity for scavenging fatty acids from LPC, as reflected in a substantially reduced reinvasion rate (Liu et al., 2024). Essentially identical results were observed here for the QKO/PARE KD line in the presence of aTc (**Fig. 3C**). Interestingly, knockdown of PARE essentially abolished the ability of parasites to complete a replication cycle in 2LPC medium (**Fig. 3C**). In contrast, deletion of PARE on a wild-type background had no effect on growth in 2LPC, which is not surprising given the high level of lysophospholipase activity provided by XL2 and XLH4 (Liu et al., 2024).

To determine where the block in development occurs for QKO/PARE KD parasites in 2LPC media, tightly synchronized cultures of wild-type (3D7) and QKO/PARE KD parasites (without and with aTc) comprising 0-4 h rings were generated using ML10 (see section 2.6) and washed into the respective media. After 53 hours, 3D7 parasites had progressed to late schizogony in both 2LPC and 2FFA media (**Fig. 3D**). In 2FFA medium, QKO/PARE KD parasites had developed similarly, both with and without aTc. In 2LPC medium, QKO/PARE KD +aTc parasites appeared similar to 3D7. In contrast, parasites in the -aTc culture exhibited smaller size, aberrant organization of nuclei, and dispersed hemozoin crystals (**Fig. 3D**), which are suggestive of an inability to form invasion-competent daughter merozoites and likely explain the near-total suppression of reinvasion observed in **Fig. 3C**.

### 3.4 PARE is a membrane-associated enzyme at the parasite periphery

To understand how PARE contributes to fatty acid acquisition from LPC, we localized the enzyme using a parasite line expressing a fusion of endogenous PARE to yellow fluorescent protein (YFP) (Elahi et al., 2019). Images of live parasites revealed that PARE-YFP accumulates at the parasite periphery at all stages of asexual development, as well as at the food vacuole membrane in trophozoites (**Fig. 4A**), which suggests that the enzyme associates with membranes. To assess whether the enzyme could be exported as a soluble protein to the parasitophorous vacuole (PV), parasites were imaged following saponin treatment, which permeabilizes the erythrocyte and PV membranes, resulting in the loss of soluble proteins. The peripheral distribution of PARE-YFP was retained in saponin-treated parasites (**Fig. 4B**), indicating that it is not primarily a soluble PV protein. To determine whether native (*i.e.*, untagged) PARE is a membrane-associated protein, a lysate of 3D7 parasites was labeled with TAMRA-FP and fractionated into soluble and membrane fractions. Nearly all labeled PARE was located in the membrane fraction (**Fig. 4C**).

**Figure 4:**
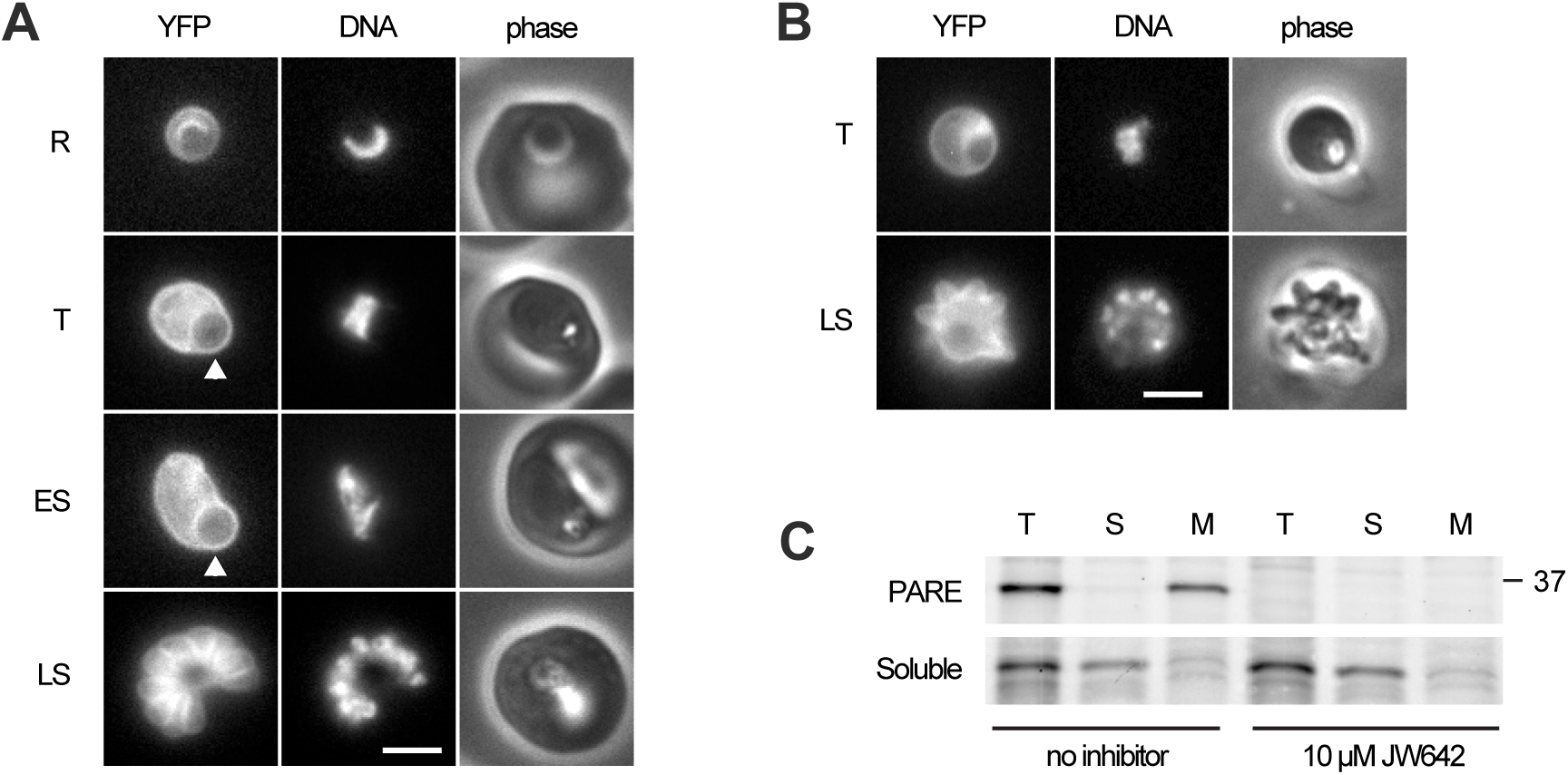
PARE is a membrane-associated enzyme. (**A**) Live parasites expressing endogenous PARE tagged with YFP at various stages of asexual development. The food vacuole is indicated with an arrowhead. (**B**) Live, saponin-treated parasites expressing PARE-YFP. In A and B: R, ring; T, trophozoite; ES, early schizont; LS, late schizont; scale bar, 3 µm. (**C**) Fractionation of lysates of wild-type 3D7 parasites that have been labeled with the serine hydrolase probe TAMRA-FP, without and with pre-treatment with the PARE inhibitor JW642 (Elahi et al., 2019). T, total lysate; S, soluble fraction; M, membrane fraction. Recovery of an unidentified serine hydrolase (estimated molecular mass ∼30 kDa) in the soluble fraction is shown to validate the separation of soluble and membrane proteins.

## 4. DISCUSSION

We set out to understand how a *P. falciparum* line that is deficient in XL/XLH lysophospholipases is able to continue to scavenge fatty acids from LPC, albeit at a reduced level that does not support robust growth (Liu et al., 2024). Our studies reveal that PARE contributes to LPC metabolism, presumably through the A-type lysophospholipase activity that is observed in the XL/XLH enzymes and other lysophospholipases of the serine hydrolase family (Liu et al., 2024; Wang and Dennis, 1999), although this remains to be confirmed. The loss of PARE had no effect on LPC hydrolysis in parasites expressing the two dominant asexual-stage lysophospholipases, XL2 and XLH4. However, when these enzymes were absent, as is the case in the QKO line, PARE-dependent lysophospholipid catabolism can be inferred from the choline-complemented, slow-growth phenotype and the reduced *in situ* LPC competition with a fluorescent fatty acid probe.

Knockdown of PARE on a QKO background impaired the parasite’s ability to scavenge essential metabolites from LPC. One of these metabolites is choline: loss of PARE elicited a reduced-growth phenotype when grown in RPMI (which contains 21 µM choline) that was complemented by supplementing the medium with 2 mM choline. This finding implies that LPC is an important source of choline even when exogenous free choline is available. It aligns with a reportthat the enzyme GDPD, which generates free choline from the LPC hydrolysis product glycerophosphocholine, is essential under standard culture conditions but not in medium supplemented with elevated levels of choline (Ramaprasad et al., 2022).

Another class of metabolites acquired from LPC hydrolysis is free fatty acids. In QKO parasites, knockdown of PARE reduced the parasite’s ability to scavenge fatty acids from LPC. The consequence of losing XL2, XLH4 and PARE activities was a near-total failure to replicate when cultured in defined media with LPC as the sole source of exogenous fatty acids. While lysophospholipase activities have been reported in erythrocytes (Podolski et al., 1983; Selle et al., 1993; Tamura et al., 1985), these are clearly insufficiently active, or are not appropriately positioned in the infected erythrocyte, to overcome the loss of the three parasite enzymes.

QKO/PARE KD -aTc cultures did progress to an early stage of schizogony, however, suggesting that a very low fatty acid-generating activity remains.

We found that PARE is a membrane-associated enzyme that accumulates at the parasite periphery and in internal structures, most notably the food vacuole. Given the apparent absence of a signal peptide or transmembrane domain, the most likely location of PARE is on the cytosolic side of the plasma membrane. This claim is supported by the retention of peripheral fluorescence in saponin-isolated parasites, which have permeabilized RBC and PV membranes. Association with the PV-facing side of the plasma membrane, or the PVM itself, cannot be ruled out; however, these possibilities seem unlikely in the absence of an export signal. Interestingly, PARE appears in the asexual *P. falciparum* “palmitoylome” (Jones et al., 2012) and therefore may be anchored to the membrane by S-acylation. Whatever its orientation, we propose that PARE catalyzes the hydrolysis of exogenous LPC that diffuses toward the parasite.

The physiological role of PARE remains enigmatic. Judging from the intensity of TAMRA-FP labeling, it is a relatively abundant parasite-encoded serine hydrolase, the loss of which does not manifest as a growth phenotype in standard culture media. We speculate that the lysophospholipase activity inferred from our studies is not physiologically relevant; rather, that PARE is a promiscuous esterase with weak lysophospholipase activity that becomes observable in an XL/XLH-deficient background. It is possible that PARE’s catalytic activity has a role to play in the maintenance of plasma membrane integrity in the host environment that does not become manifest under culture conditions. Alternately, it may have a function in another of the parasite’s life cycle stages. Resolving these different possibilities will require further study.

## CRediT AUTHORSHIP CONTRIBUTION STATEMENT

**Jiapeng Liu**: Conceptualization, methodology, investigation, writing-review & editing, visualization.

**Katie Fike**: Methodology, investigation.

**Seema Dalal**: Methodology, investigation.

**Christie Dapper**: Methodology, investigation.

**Michael Klemba**: Conceptualization, funding acquisition, methodology, validation, writing-original draft, visualization, supervision, project administration.

## ACKNOWLEDGMENTS

We are grateful to D. Goldberg for the parasite line 3D7-DR4, J. Beck for CRISPR plasmids, S. Prigge for the TetR-DOZI plasmid, T. Nevalainen for AKU-010, and Christiaan van Ooij and LifeArc for ML10.

