## Supplementary Table 1 for "Defining the minimal enzymatic requirements for fatty acid scavenging from lysophosphatidylcholine by erythrocytic *Plasmodium falciparum*"

**Supplementary Table 1: Oligonucleotides used in this study.**

| Oligo | Location | Function | Sequence |
|---|---|---|---|
| 1381 | PARE CDS | sgRNA | TAAGTATATAATATTTACTTGTTCTTCTTG TTTGGGTTTTAGAGCTAGAA |
| 1382 | PARE CDS | sgRNA | TTCTAGCTCTAAAACCCAAACAAGAAGA ACAAGTAAATATTATATACTTA |
| 1387 | PARE CDS | HA1, repair plasmid | GATATCGTCCACCTGGATATCTTTACGTC TTACTCCTGGTTTACG |
| 1388 | PARE CDS | HA1, repair plasmid | CATAAGGATAGACGTCTACTTGTTCTTCC TGCTTAGGGGTATGGACAGCTAGCCATG* |
| 1389 | PARE 3’ UTR | HA2, repair plasmid | CCCTTTCCGGGCGCGCCTTTAATGAAGCG TATTATACTGACG |
| 1390 | PARE 3’ UTR | HA2, repair plasmid | GATATCCAGGTGGACGATATCGGAAGAA TGGATTAAGAGAAAAAG |
| 1413 | PARE CDS | Genotyping PCR | AGCTGGTATGATATCTATAGATGAG |
| 1407 | pKD-PARE | Genotyping PCR | CCAGACAGTGGCCCTTATGCG |
| 1214 | pKD-PARE | Genotyping PCR | ATTATTATATTTAACTATATACTATGG |
| 1414 | PARE 3’ UTR | Genotyping PCR | CAATTTTGTATACACAAAATAAGTTGC |

Abbreviations: CDS, coding sequence; HA, homology arm; UTR, untranslated region.

* Shield mutations are indicated in red.
